## Supplementary Information for "Structure and dynamics of Toll immunoreceptor activation in the mosquito Aedes aegypti"

**List of Supplementary Figures**

- **Supplementary Figure 1: Cryo-EM workflow.**
- **Supplementary Figure 2: Effect of density modification on cryo-EM maps.**
- **Supplementary Figure 3: Comparison of Aedes and Drosophila Toll and Spz structures.**
- **Supplementary Figure 4: Asymmetry in structure and sequence variation in Spz.**
- **Supplementary Figure 5: Shape and electrostatic complementarity of Toll5A and Spz1C.**
- **Supplementary Figure 6: Spz1C binding to Toll5A Z-loop.**
- **Supplementary Figure 7: In-gel mass fingerprinting of Toll5A and its degradation product.**
- **Supplementary Figure 8: Surface plasmon resonance suggests paralog and orthologue specificity.**
- **Supplementary Figure 9: Solution scattering data of Toll5A in the absence and presence of Spz1C.**
- **Supplementary Figure 10: Analytical ultracentrifugation profiles.**
- **Supplementary Figure 11: Aag2 transcriptional profile of members of the Toll and Spaetzle family.**
- **Supplementary Figure 12: Bioinformatic analysis.**

**List of Supplementary Tables**

- **Supplementary Table 1: Cryo-EM Data Collection, refinement, and validation Statistics.**
- **Supplementary Table 2: Solution scattering of Toll5A alone and in complex with Spz1C**
- **Supplementary Table 3: RT-PCR oligonucleotides used.**

**List of Supplementary Source data**

- **Supplementary Source data 1: Uncropped gels relating to Supplementary Fig. 6b.**
- **Supplementary Source data 2: Uncropped gel relating to Supplementary Fig. 7.**
- **Supplementary Source data 3: Agarose gels relating to Supplementary Fig. 11.**

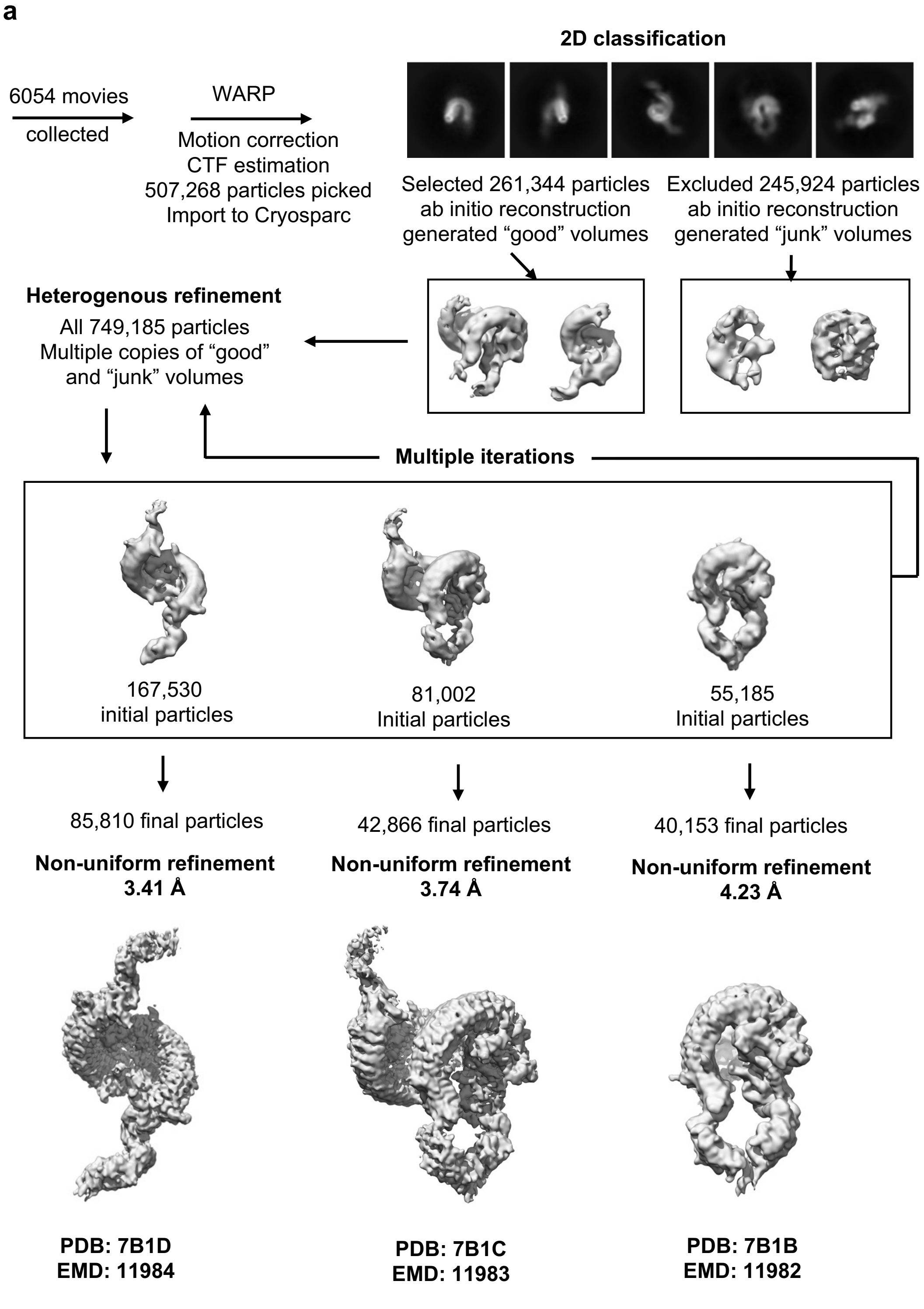
a
2D classification
6054 movies
collected
WARP
Motion correction
CTF estimation
507,268 particles picked
Import to Cryosparc
Selected 261,344 particles
ab initio reconstruction
generated “good” volumes
Excluded 245,924 particles
ab initio reconstruction
generated “junk” volumes
Heterogenous refinement
All 749,185 particles
Multiple copies of “good”
and “junk” volumes
Multiple iterations
167,530
initial particles
81,002
Initial particles
55,185
Initial particles
85,810 final particles
Non-uniform refinement
3.41 Å
42,866 final particles
Non-uniform refinement
3.74 Å
40,153 final particles
Non-uniform refinement
4.23 Å
PDB: 7B1D
EMD: 11984
PDB: 7B1C
EMD: 11983
PDB: 7B1B
EMD: 11982

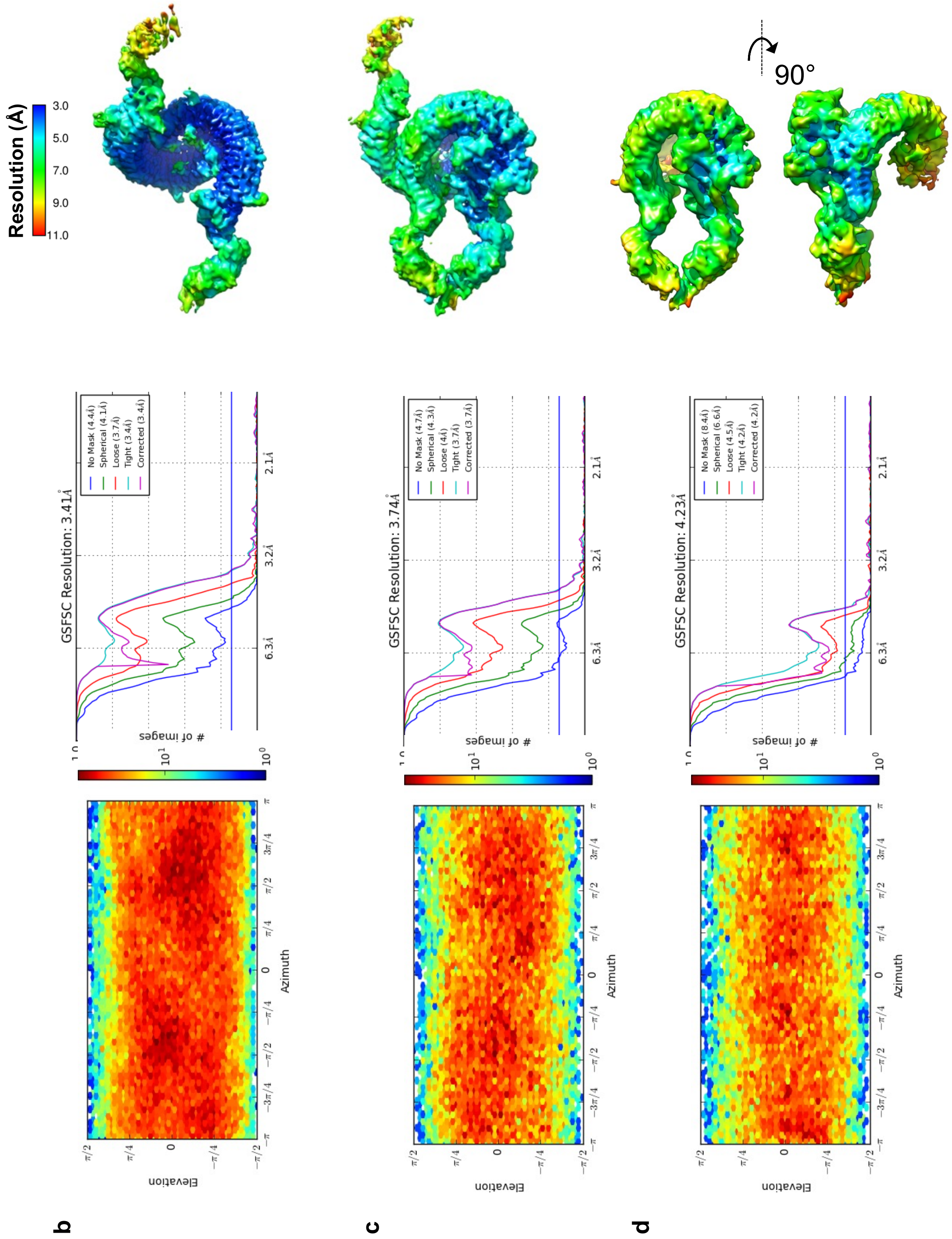

<u>**Supplementary Figure 1**</u>**: Cryo-EM workflow. a** Data were collected on a Titan Krios. Representative cryo-EM micrographs are shown. Scale bars, 50 nm. Cryo-EM data analysis was performed in Warp[1] and CryoSparc[2]**.** FSC threshold criteria at 0.143[3] is used to determine the resolution of the reconstructions and indicates the size of the smallest reliable detail. The atomic model refinement process was halted when iterations stopped improving. In the end, the maps were masked, sharpened, and a final resolution was determined. **b** Toll5A homodimer, **c** Spz1C-bound Toll5A trimer. **d** Spz1C-bound Toll5A heterodimer. For each particle the following are shown: First column: the angular distributions for particles projections estimated by cryoSPARC[2]. Heat maps show number of particles for each viewing angle from low, in dark blue, to high, in red. Second column: the gold standard FSC, calculated by comparing the two independently determined half maps from cryoSPARC. The purple line represents the 0.143 FSC cut-off, which indicates a nominal resolution after correction. Third column: density coloured according to a local resolution gradient from high at 3 Å, in dark blue; to low resolution, with 7 Å in green, and 11 Å in red.

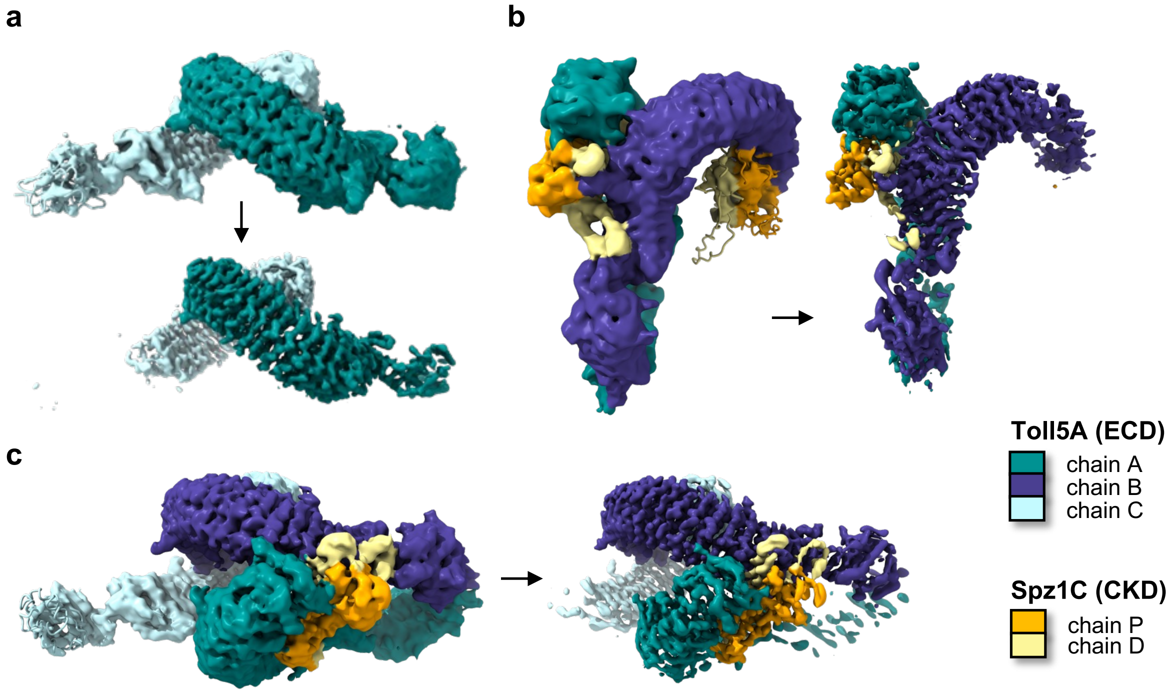

__Supplementary Figure 2__: **Effect of density modification on cryo-EM maps. a** Homodimer before and after density modification[4]. **b** Heterodimer. **c** Heterotrimer. Maps are coloured according to protein content. Figure generated in ChimeraX (https://www.rbvi.ucsf.edu/chimerax)[5].

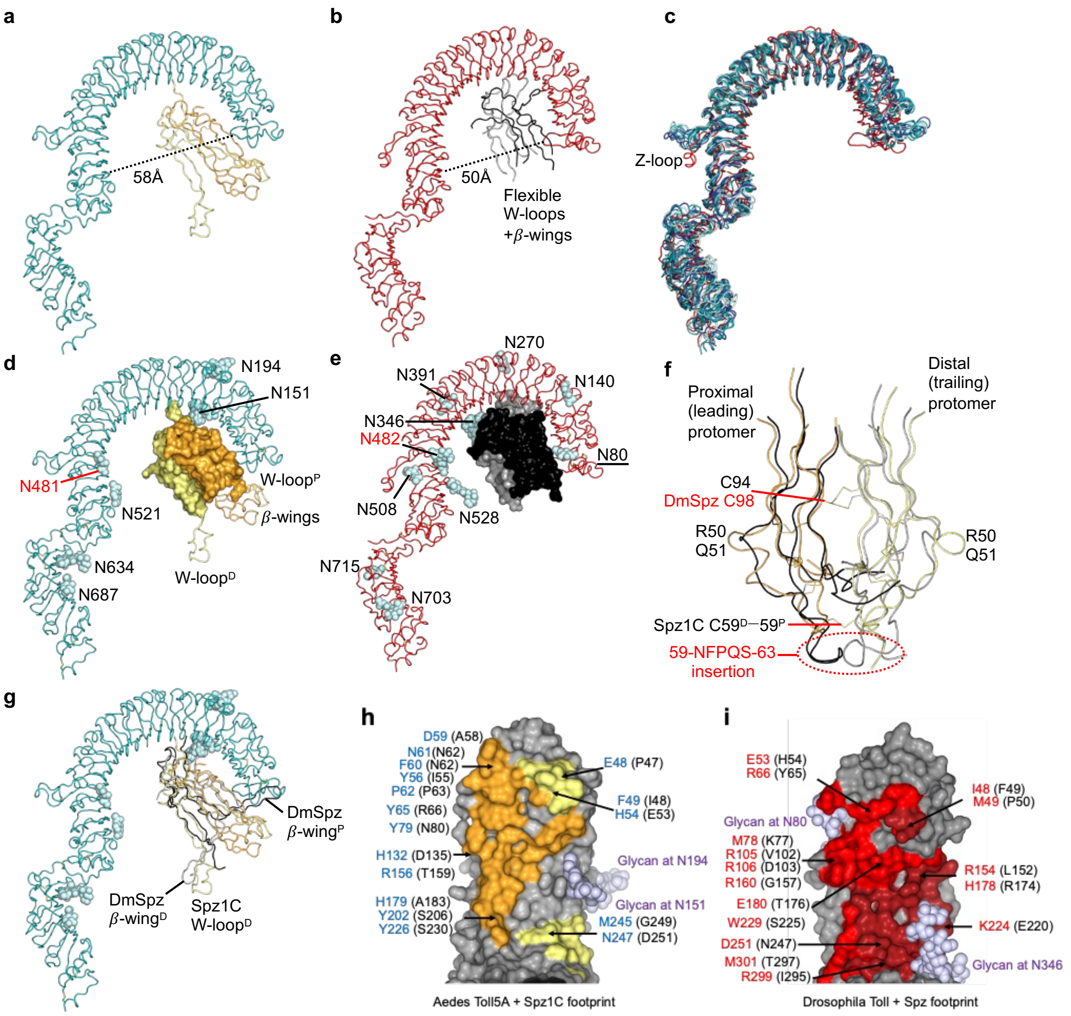

**Supplementary Figure 3: Comparison of Aedes and Drosophila Toll and Spz structures.** Automatic secondary structure matching (SSM)[6] was used to superpose the equivalent secondary structural elements of proteins. **a** Aedes Spz1C-ligated Toll5A chain A, as observed in the heterodimeric complex. **b** Drosophila Toll bound to Spz[7]. **c** SSM-generated superposition of Drosophila Toll and Toll5A chains from homodimers (chains A and C), heterodimers (chains A and B) and heterotrimers (chains A, B and C). **d** Toll5A has six visible Asn-linked glycosylation sites. Ribbon diagram representation for Toll5A, blue-white spheres for glycosylations, partial molecular surface of Spz1C with Trp-loops and $\beta$-wings shown as ribbons, for comparison with Drosophila. Asn-151 glycans are located on the descending flank of LRR2, within 10 Å of the distal C-terminal region of Spz1C. Labelled in red, Asn-481 (ascending LRR15) in Aedes Toll5A, and Asn-482 in Drosophila, respectively, are the only conserved glycosylation sites. **e** Drosophila Toll has thirteen visible Asn-linked glycosylation sites. Glycosylation at Asn-80 prevent Drosophila from adopting a mosquito-like binding mode. **f** Close-up on a Spz cys-knot core region overlay. Disulphide bonds are shown in lines. Cys-59 in Aedes Spz1C, which displays an additional intermolecular disulphide bond, caps off an area of deletion compared to other Spz isoforms (residues 59 to 63 in Drosophila Spz). In Drosophila, the corresponding insertion provides extensive contacts at the primary Toll binding side. Aedes Spz1C has a unique insertion with Arg-50 and Gln-51 that forms a small protrusion, which interacts with $Asn521^{B}$-linked glycans at the dimer interface. **g** Overlay of refolded Drosophila Spz[8] with the mosquito structure. The perpendicular distal $\beta$-wing of Drosophila is reminiscent of the extended conformation of the mosquito distal Trp-loop. **h** Spz1C footprint on Toll5A at the concave binding site. Molecular surface of the N-terminal region of Toll5A, in grey. Contact residues are given for Toll5A, with their Drosophila counterpart in brackets. **i** Drosophila Spz footprint on Toll. Molecular surface of the N-terminal region of Toll, in grey. Contact residues are given for Toll, with their Toll5A counterpart in brackets. Structural differences, glycosylation, along the divergence in primary sequences, help rationalise the lack of interaction of Spz isoforms other than Spz1C with Toll5A.

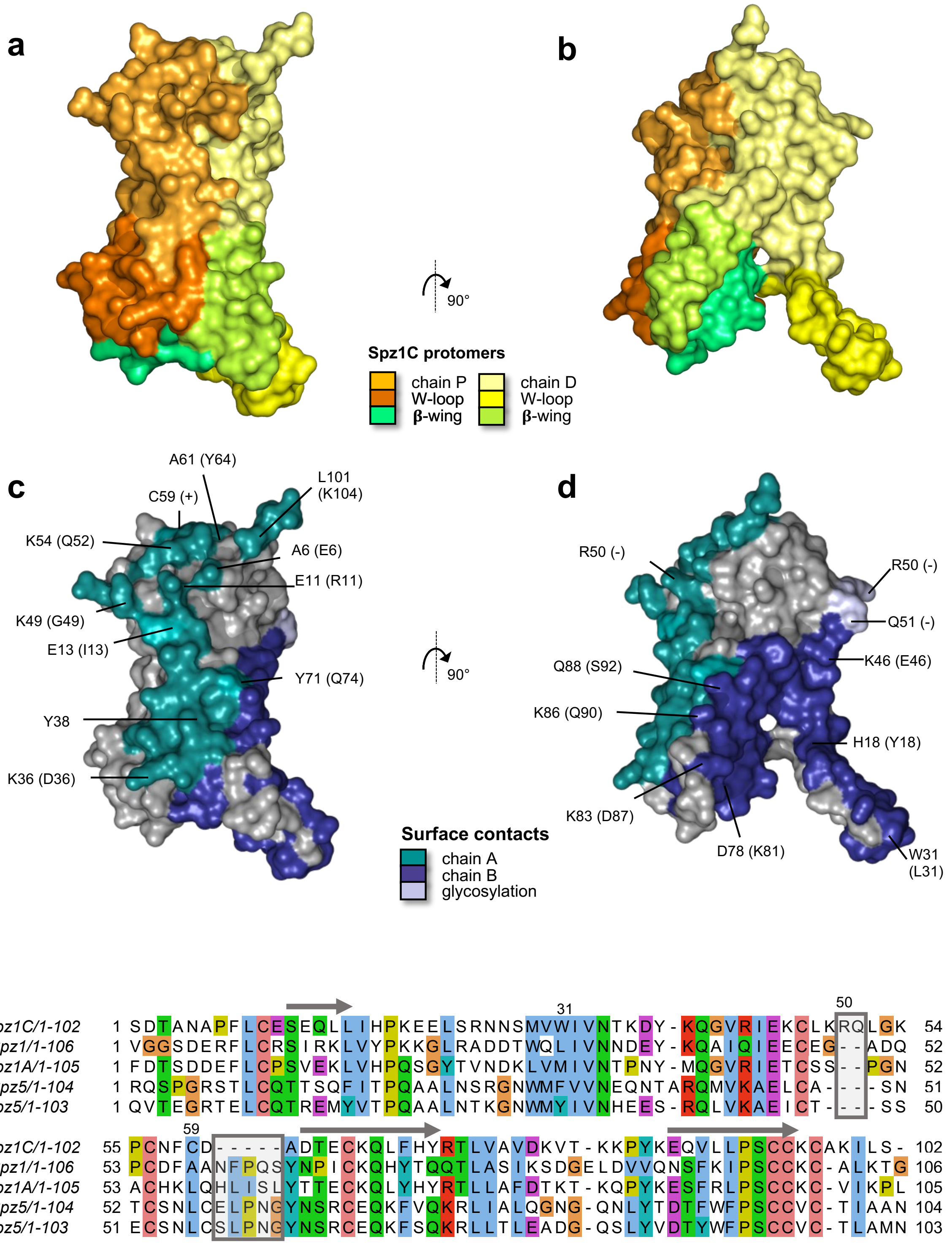

<u>**Supplementary Figure 4:**</u> **Asymmetry in structure and sequence variation in Spz.**
**a-b** Molecular surfaces of Spz1C in two top views rotated by 90° and color-coded according to each protomer. **c-d** Molecular footprint of Toll5A on Spz1C. The corresponding residue in Drosophila Spz is indicated in brackets to highlight sequence variation in the molecule. **e** Sequence alignment. *β*-strands are indicated by a grey arrow above the sequence, main insertion and deletion compared to Drosophila are boxed.

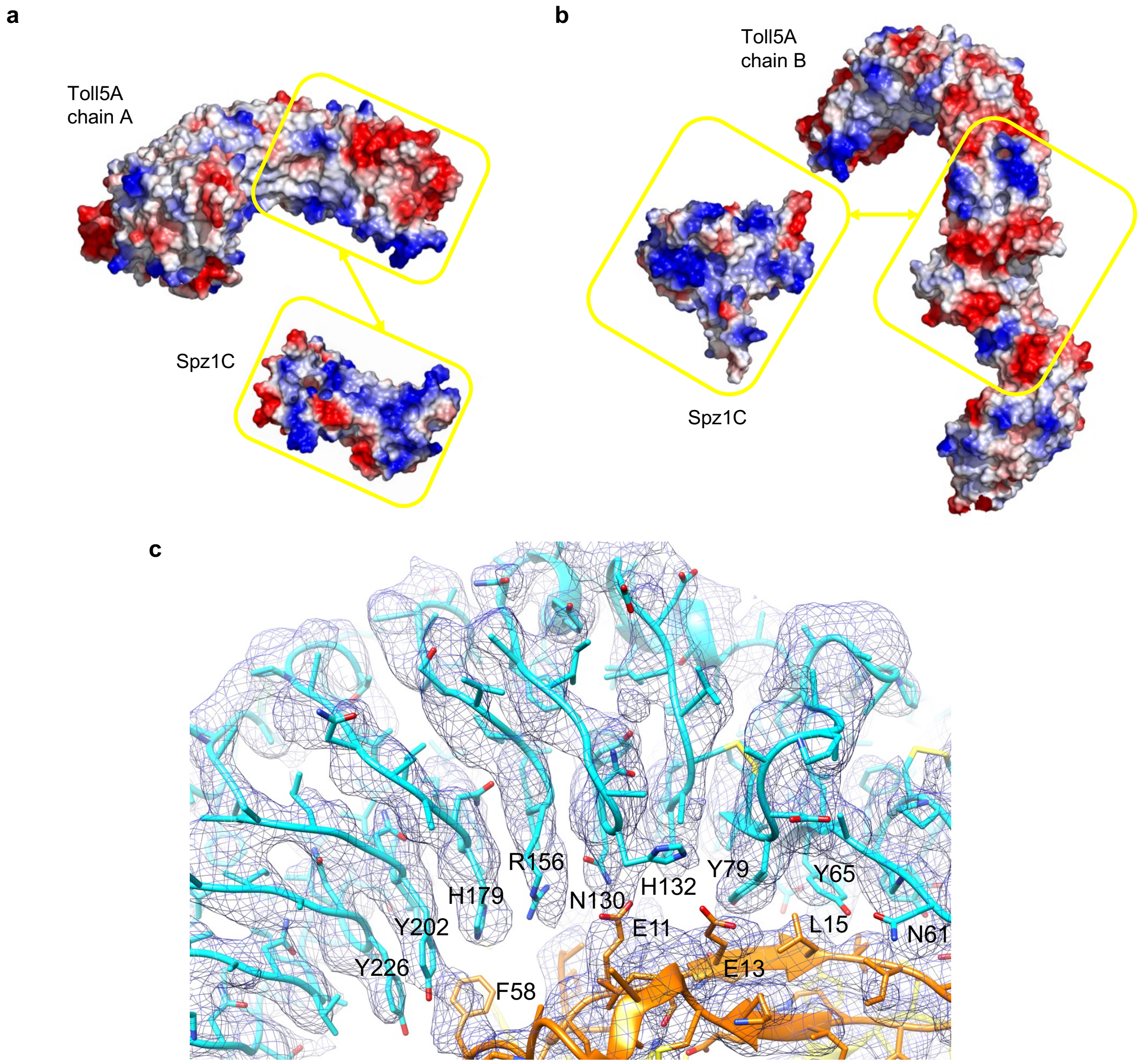

**Supplementary Figure 5: Shape and electrostatic complementarity of Toll5A and Spz1C.**
All-atom models were generated in Modeller[9]. Molecular surface charge distributions were generated in PyMol (https://pymol.org/) with electronegative patches in red, neutral in white and electropositive in blue. Focusing on the concave binding site in **(a)** looking up towards Toll5A and down onto Spz1C interacting surface representations. Focusing on the convex lateral interface in **(b)** with complementary views that reveal Spz1C and Toll5A interfaces. **(c)** Close-up cartoon representation with density map (blue wire) of the concave interface.

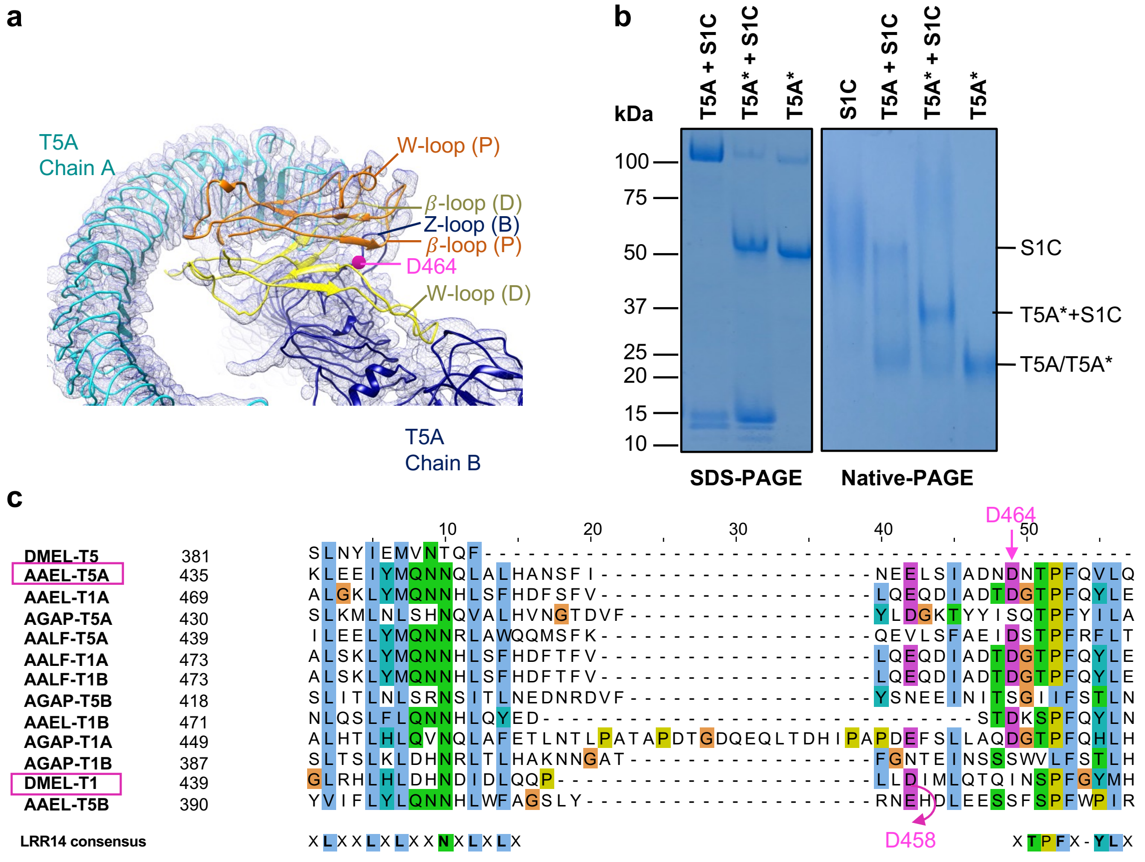

**Supplementary Figure 6: Spz1C binding to Toll5A Z-loop.** Electron density of unprocessed Toll5A-Spz1C 2:1 heterodimer with Spz1C crosslinking the Z-loop at the heterodimerization interface. Spz1C $\beta$-loops surround the Z-loop, while the W-loops connect the N-terminal region of chain A and the hinge region of chain B, respectively. The location of Toll5A Asp 464 is indicated by a magenta sphere (**a**). Spz1C protects Toll5A ECD from spontaneous degradation (purified sample stored at 4°C for 3 weeks). In contrast, spontaneous Z-loop degradation stabilises Spz1C binding with a clear band shift as evidenced by native gel electrophoresis (**b**). Sequence alignment of duplicated mosquito Tolls and *Drosophila* Toll1 and 5 at LRR14 and its poorly conserved Z-loop (**c**). DmToll5 (DMEL-T5) does not have a Z-loop. *A. aegypti* Toll5A (AAEL-T5A) Asp 464 (highlighted in magenta) is partially conserved in most mosquito receptors and a potential target site for Asparaginyl endopeptidase (AEP) procession. An alternative location is described for *Drosophila* Toll at Asp 458. Some receptors have an Asn substitution instead of Asp, as for *A. gambiae* Toll1B and 5B, while the site's position does not appear conserved for *A. aegypti* Toll5B, with its Asn at +1 a position and Asp at +3. *Drosophila* Toll Asp 458 cleavage is involved in constitutive receptor dimerization[10]. Sequence alignment generated by Muscle[11] software and illustrated with ClustalX colour scheme above 30 % conservation in Jalview[12]. *A. albopictus* AALF-T1A (AALF013620); AALF-T1B (AALF013620); AALF-T5A (AALF027669); *A. aegypti* AAEL-T1A (AAEL026297) ; AAEL-T1B (AAEL027951) ; AAEL-T5A (AAEL007619, Uniprot A0A6I8TEX2); AAEL-T5B (AAEL000057) ; *A. gambiae* AGAP-T1A (AGAP001004 ; UniProt Q8WRE5) ; AGAP-T1B (AGAP010636; Q7QE54) ; AGAP-T5A (Q5TW82 ; AGAP000999) ; AGAP-T5B (Q7QE40 ; AGAP010669) ; *D. melanogaster* Toll1 (DMEL-T1: UniProt P08953) and Toll5 (Q9NBK8).

**Fixed modifications:** Carbamidomethyl (C)
**Variable modifications:** Deamidated (NQ), Oxidation (M)

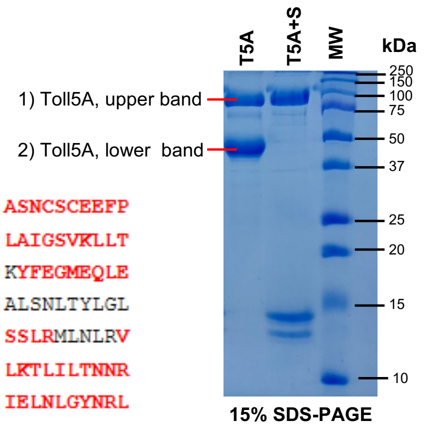

15% SDS-PAGE

**Protein sequence coverage: 56%**

### 1) Toll5A, upper band

Matched peptides shown in ***bold red***.

```
   1 MNRTHRSIAL LVISAIILLV QTQDVTSTST KRFTCPEESE ASNCSCEEFP
  51 SKTHFYCPDF NPTLYVDVED RMRVDFKCYD EPHDFKSLPN LAIGSVKLLT
 101 VVDCVLDDDR PILESFKFLE VADVRSFVYN NHENGIRYNA KYFEGMEQLE
 151 NLTLARGVVS IDRDTFSGFL NLKRLTIEHN KLNLQPGTFE ALSNLTYLGL
 201 VYNGLNEIQP GLFDGLESLE ALSLSYNDIK SLSAGSFNGL SSLRMLNLRV
 251 NKIESFDANT FASLKELSRL EITLNPFVSL PRGLFSENKK LKTLILTNNR
 301 KLVTLPEELL ANLKELTVVN LSHNGVGNLP ESLLSGSSGI IELNLGYNRL
 351 NSLPEELLSD QPQLQVLNLD HNQLESIPDY FLERNVELQT LYLSHNRLRS
 401 LSEKAFTKLK NLKELHLENN QLQTIPQFLF SGTPKLEEIY MQNNQLALHA
 451 NSFINEELSI ADNDNTPFQV LQKLRILHLR NNSISTIFQD WYINNLEMQS
 501 LDLSFNKLPG LSYTQLQFQS NITLNLSNNE ISQVLLIDDL DLQPYQRINV
 551 DLNHNPLNCN CNALKFIQLI QSKAEHGLQF NVDQLRCSEP PNLLDATMDQ
 601 LQTKDLLCDF ESADDCPKDC QCAMRLLDHT VIVNCSGRGL TEFPDLPIPS
 651 QLHEDFNALE VHVENNRLTK LPNLTKHNEI TQLYARNNSI QNLLPHNIPS
 701 KLRIIDLSQN LLKMIDDSTL AQINRSSHLE TIRLSQNQWL CDCPASSFLI
 751 FVQQNSRLIS DMSAIRCHPS GKSLDSITVN ELCFEDYTTK IVICFAIAVF
 801 GLLVGLISLL FFRYQTEVKV WLFTHNLFLS LITEEELDKD KLYDAFISYS
 851 HLDEEFIVDE LIPKLENDPM NFKTCWHVRD FMPGEMIMTQ IVKSIEASRR
 901 TVIVLSKNFL ESSWAKQEFR QAHVQSMEDN RVRVIVVIYE DIGDIDSLDG
 951 ELKAYLKTNT YVKWGDPWFW QKLRYAMPHP LKVKGIKSAV SEIHLQQVTT
1001 KATPTAV
```

**Protein sequence coverage: 52%**

### 2) Toll5A, lower band

Matched peptides shown in ***bold red***.

```
   1 MNRTHRSIAL LVISAIILLV QTQDVTSTST KRFTCPEESE ASNCSCEEFP
  51 SKTHFYCPDF NPTLYVDVED RMRVDFKCYD EPHDFKSLPN LAIGSVKLLT
 101 VVDCVLDDDR PILESFKFLE VADVRSFVYN NHENGIRYNA KYFEGMEQLE
 151 NLTLARGVVS IDRDTFSGFL NLKRLTIEHN KLNLQPGTFE ALSNLTYLGL
 201 VYNGLNEIQP GLFDGLESLE ALSLSYNDIK SLSAGSFNGL SSLRMLNLRV
 251 NKIESFDANT FASLKELSRL EITLNPFVSL PRGLFSENKK LKTLILTNNR
 301 KLVTLPEELL ANLKELTVVN LSHNGVGNLP ESLLSGSSGI IELNLGYNRL
 351 NSLPEELLSD QPQLQVLNLD HNQLESIPDY FLERNVELQT LYLSHNRLRS
 401 LSEKAFTKLK NLKELHLENN QLQTIPQFLF SGTPKLEEIY MQNNQLALHA
 451 NSFINEELSI ADNDNTPFQV LQKLRILHLR NNSISTIFQD WYINNLEMQS
 501 LDLSFNKLPG LSYTQLQFQS NITLNLSNNE ISQVLLIDDL DLQPYQRINV
 551 DLNHNPLNCN CNALKFIQLI QSKAEHGLQF NVDQLRCSEP PNLLDATMDQ
 601 LQTKDLLCDF ESADDCPKDC QCAMRLLDHT VIVNCSGRGL TEFPDLPIPS
 651 QLHEDFNALE VHVENNRLTK LPNLTKHNEI TQLYARNNSI QNLLPHNIPS
 701 KLRIIDLSQN LLKMIDDSTL AQINRSSHLE TIRLSQNQWL CDCPASSFLI
 751 FVQQNSRLIS DMSAIRCHPS GKSLDSITVN ELCFEDYTTK IVICFAIAVF
 801 GLLVGLISLL FFRYQTEVKV WLFTHNLFLS LITEEELDKD KLYDAFISYS
 851 HLDEEFIVDE LIPKLENDPM NFKTCWHVRD FMPGEMIMTQ IVKSIEASRR
 901 TVIVLSKNFL ESSWAKQEFR QAHVQSMEDN RVRVIVVIYE DIGDIDSLDG
 951 ELKAYLKTNT YVKWGDPWFW QKLRYAMPHP LKVKGIKSAV SEIHLQQVTT
1001 KATPTAV
```

Missing peptide in Toll5A, lower band

**Supplementary Figure 7: In-gel mass fingerprinting of Toll5A and its degradation product.**
During peptide mass fingerprinting the target proteins were digested with trypsin to yield the constituent small peptides. The accurate mass of these peptides is determined by MS analysis. This gives the peak list of peptides (in bold red). This peak list is compared with the theoretical peptide peak list obtained from the *in silico* digestion of Vectorbase database[13] and the best match is identified by computer software. A similar degradation pattern was reproduced using asparagine endopeptidase AspN from Flavobacterium menigosepticum (NEB Cat. No. P8104S).

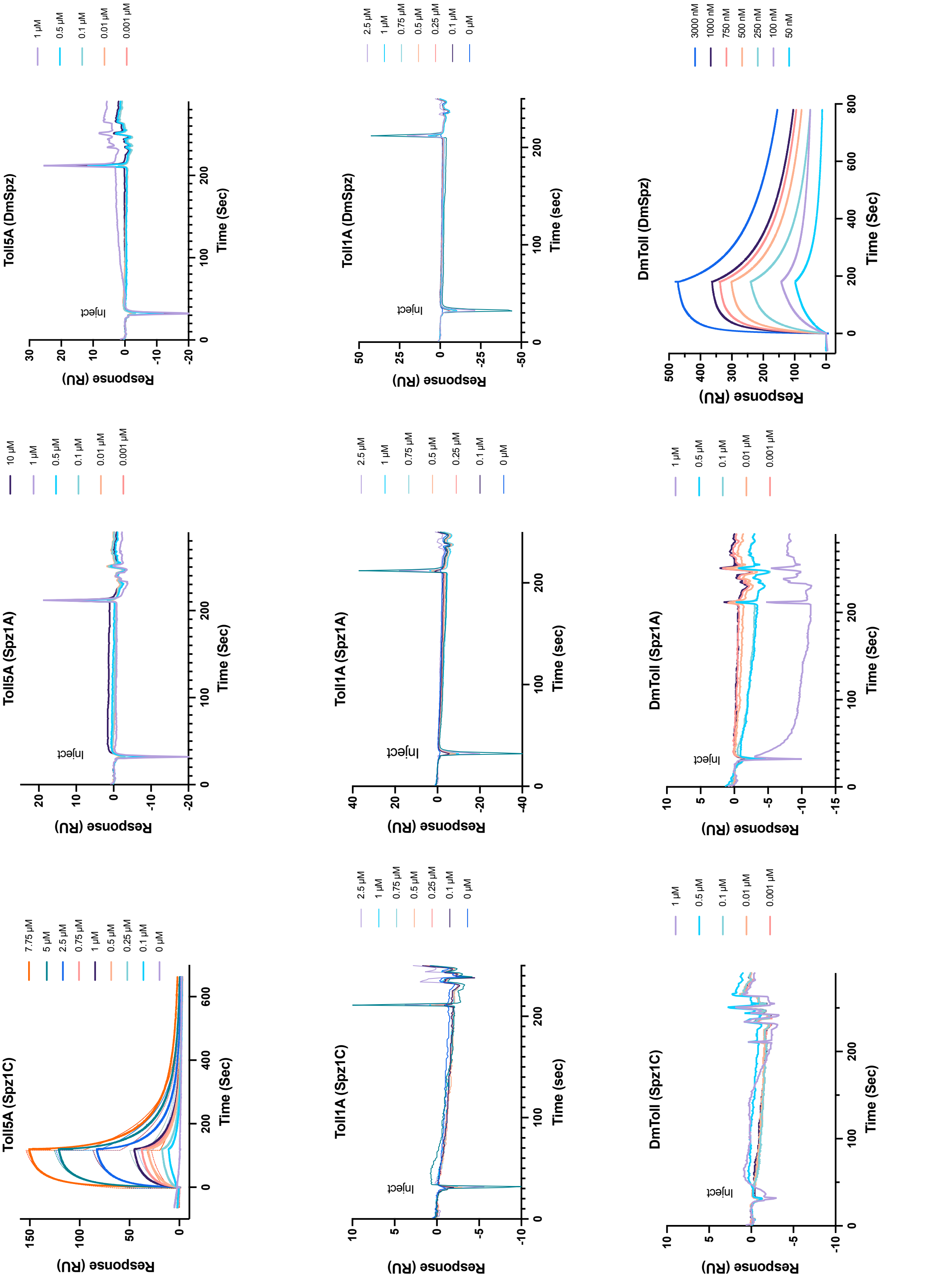

Toll5A (Spz1C)
Toll5A (Spz1A)
Toll5A (DmSpz)
Toll1A (Spz1C)
Toll1A (Spz1A)
Toll1A (DmSpz)
DmToll (Spz1C)
DmToll (Spz1A)
DmToll (DmSpz)
Response (RU)
Time (Sec)
Time (sec)
Inject
7.75 µM
5 µM
2.5 µM
0.75 µM
1 µM
0.5 µM
0.25 µM
0.1 µM
0 µM
10 µM
0.01 µM
0.001 µM
3000 nM
1000 nM
750 nM
500 nM
250 nM
100 nM
50 nM

| Toll5A-Spz1C | $k_{on}$ (1/Ms) | $k_{off}$ (1/s) | $K_D$ ($\mu$M) | Chi[2] | $R_{max}$ (RU) |
|---|---|---|---|---|---|
| Global heterogeneous | $k_{on\,1}$ = 16,930 $k_{on\,2}$ = 2,548 | $k_{off\,1}$ = 0.03239 $k_{off\,2}$ = 0.005325 | $K_{D\,1}$ = 1.913 $K_{D\,2}$ = 2.090 | 1.65 | $R_{max\,1}$ = 93.45 $R_{max\,2}$ = 66.40 |

**Supplementary Figure 8: Surface plasmon resonance suggests paralog and orthologue specificity.** Sensorgrams and global fitting analysis of heterogeneous ligand with two binding sites of Toll5A and Spz1C. Both binding sites have similar affinities despite different *on* and *off* rate constants.

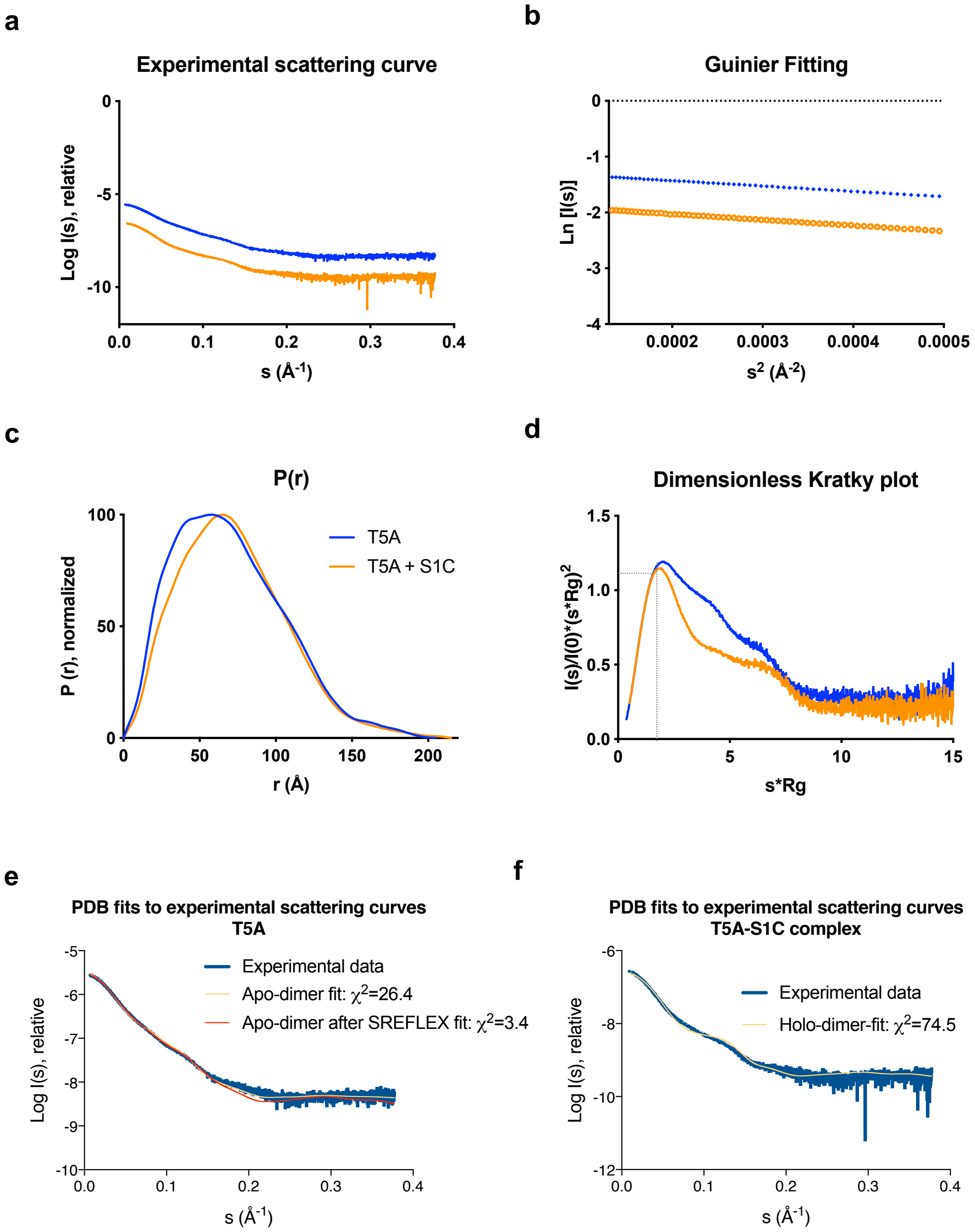

**g**

| PDB | .dat file | $R_g$ (Crysol) Å | $\chi^2$ (Crysol) | $R_g$ (Pepsi-SAXS) Å | $\chi^2$ (Pepsi-SAXS) |
|---|---|---|---|---|---|
| Apo-monomer | T5A | 48.54 | 1149.9 | 46.9 | 894.36 |
| Apo-dimer | T5A | 56.65 | 26.35 | 55.3 | 33.22 |
| Holo-dimer | T5A | 49.37 | 257.47 | 49.2 | 271.5 |
| Apo-monomer | complex | 48.54 | 2107.23 | 46.9 | 1814.4 |
| Apo-dimer | complex | 56.81 | 217.68 | 55.3 | 185.2 |
| Holo-dimer | complex | 50.43 | 74.51 | 49.2 | 68.9 |

**Supplementary Figure 9: Solution scattering data of Toll5A in the absence and presence of Spz1C. a** 1-dimensional intensity plot with **b** its Guinier fit. **c** Normalized Pair-distance distribution function, used to determine the maximum dimension of the particles ($D_{max}$). **d** Dimensionless Kratky Plot, which quantifies the conformational flexibility of proteins. The dotted lines are drawn at $qRg = 1.73$ and $I(q)/I(0)\cdot(qRg)^2=1.1$. Folded proteins have a local maximum where the two lines intersect [14]. **e-g** PDB structures were fitted to experimental curves. Dimers are the best fit using Crysol in ATSAS version 2.8.3[15] and Pepsi-SAXS version 2.6[16]. Using SREFLEX (flexible refinement of high-resolution models based on SAXS and normal mode analysis, ATSAS version 2.8.3)[17], the revised value of $\chi^2$ (Crysol) for the apo-dimer modified by SREFLEX = 3.4. However, SREFLEX did not work with the heterodimer complex. Predictions with the program OLIGOMER[18] were carried out to check if a combination of dimer-monomer worked better as a fit to the Toll5A data set. As suggested by SEC-SAXS, this was not the case, and the best fit remains for the dimer models.

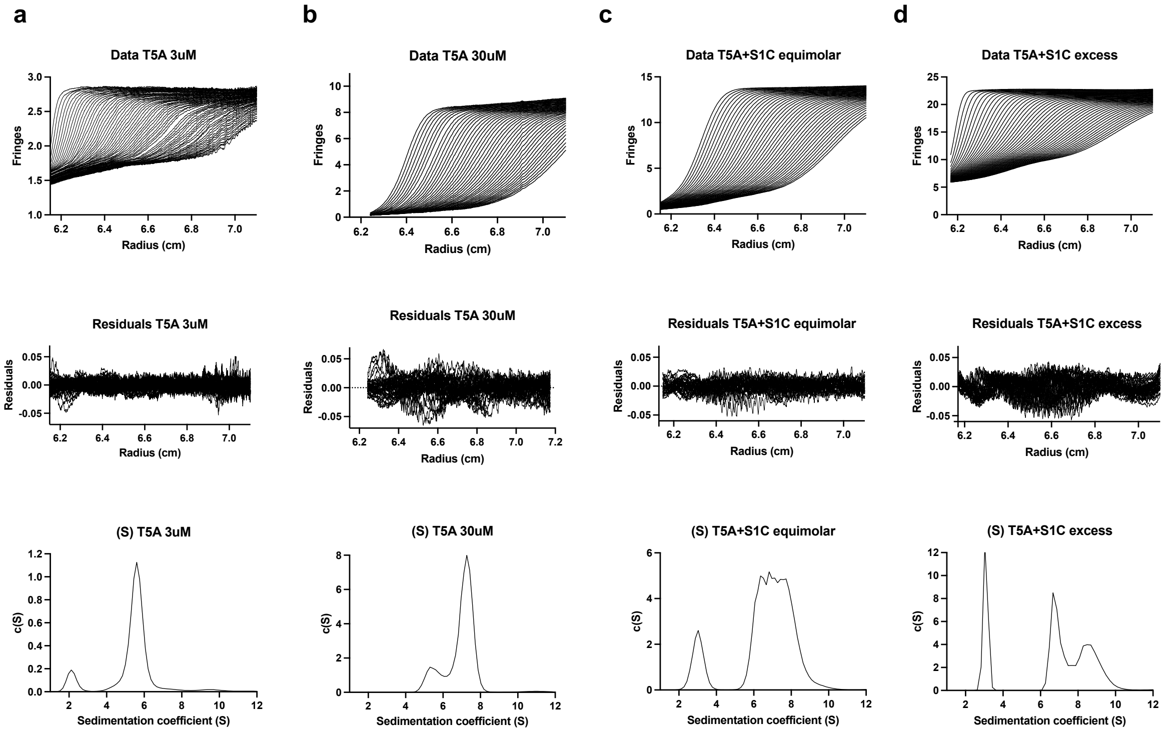

**Supplementary Figure 10: Analytical ultracentrifugation profiles.** AUC sedimentation data are shown with their fit and residuals after fitting to a c(S) model in SEDFIT and the distribution of sedimentation coefficients. From left to right: **a** 3 $\mu$M Toll5A; **b** 30 $\mu$M Toll5A; **c** mixture of 30 $\mu$M Toll5A and 30 $\mu$M Spz1C (equimolar); and **d,** mixture of 30 $\mu$M Toll5A and 74 $\mu$M Spz1C (excess). There is a contaminant at ~ 2S in the Toll5A preparation, which is only visible in the 3 $\mu$M Toll5A dataset given its lower c(S) scale.

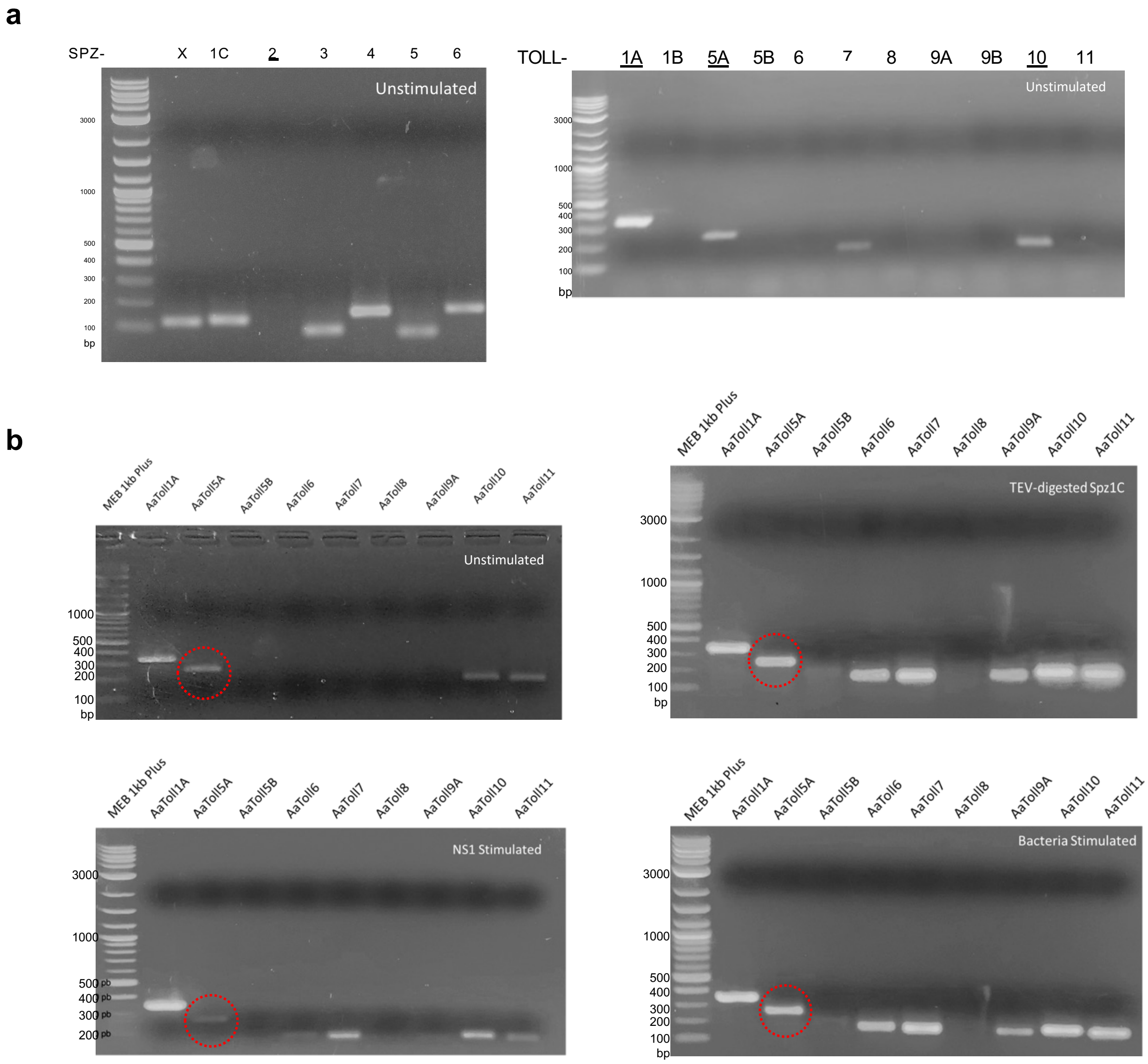

**<u>Supplementary Figure 11</u>: Aag2 cells transcriptional profile of members of the Spz and Toll family.** RT-PCR reactions were analysed on 1.5% agarose gel using SYBR™ Safe DNA gel stain in duplicates. RT-PCR primers are listed in **Supplementary Table 3**. (**a**) Unstimulated Aag2 cells express all but Spz2, along with Toll1A, Toll5A, Toll10. (**b**) Stimulated and unstimulated cells are compared for Toll expression under various conditions: addition of cleaved Spz1C (20 nM); purified Dengue virus NS1 protein (20 nM) and, heat-inactivated Gram-negative bacteria extract (5 ng/$\mu$l). Note that levels of Toll7 and Toll11 mRNA undergo batch-to-batch variations in our hands, bringing them above or below detection levels.

**Supplementary bioinformatic analysis.**
In mosquitoes, immune-related gene families involved in pattern recognition and effector activity have increased compared to *Drosophila* as a result of gene duplication and family expansion[19]. In contrast, cytosolic signal transducers such as MyD88 tend to have a stricter orthology. At the interface between both are transmembrane receptors and their ligands. Spz1 is the only ligand that has diversified in *A. aegypti* with genetic evidence pointing towards a specific role for Spz1C in mosquito immunity. In particular, Spz1C together with Toll5A mediate anti-fungal immunity[20]. More importantly, Dengue virus upregulates Spz1C in the midgut[21] and Toll5A in the salivary glands[22], hinting at a role in vector-virus interactions.

At a protein level, mosquito Toll-1 and 5 receptors are between 32 – 41 % identical to *Drosophila* Toll (DmToll-1) and between 36 to 53 % identical within the *Aedes* Toll-1/5 paralogue group. However, the extracellular domains alone are less well conserved (27 – 35 %). This contrasts with the non-duplicated receptors, which have sequence identities typically over 60 %. Upon closer inspection, *Aedes aegypti* Toll-1A, 1B and 5A have a precisely conserved number of leucine-rich repeats (LRRs) and Cys-rich capping structures, comparable to the prototypical *Drosophila* Toll-1 receptor (**Fig. S12**). On the other hand, Toll5B differs, with the gene being located on chromosome III and a shorter ectodomain with 15 LRRs instead of 17 at the N-terminus. The duplicated mosquito Toll-5 receptors also differ markedly from the *Drosophila* orthologue DmToll-5 called Tehao, which has 8 N-terminal LRRs and fewer cysteine residues. No ligand for DmToll-5 has been identified so far. We note that its ectodomain lacks features involved in DmToll-1 ligand binding. Nevertheless DmToll-5 can induce the production of antimicrobial peptides when overexpressed in cell culture and forms heterodimers with DmToll-1[23].

As for the Toll ligand, the duplicated *Spz1* genes in *Aedes* have also diverged considerably from *Drosophila* with only 21-31% sequence identity among orthologues. Non-duplicated *Spz* vary from 35 % for *Spz2* to 89 % for *Spz6*. The C-terminal active fragment containing the Cys-knot domain follows the same trend and displays sequence identities around 64-88 % for 1-to-1 orthologues, and 25-33 % for duplicated *Spz1*. The direct orthologue of *DmSpz1* is predicted to be *SpzX* in *A. aegypti*. *SpzX* also conserves the patterns of alternative splicing seen for *DmSpz1* including transcripts with a truncated Cys-knot (*SpzX-G*)[24] (**Fig. S12**). However, *SpzX* represents the amalgamation of two different Cys-knot sequences previously annotated *Spz1A* and *Spz1B*, now gathered in one locus that undergoes alternative splicing[20,25,26]. Remarkably, four splice forms *SpzX-D/E/I/J*, have Cys-knot domain sequences identical to *Spz1A* but gained an additional C-terminal furin cleavage site. *SpzX-D* and *E* acquired an additional furin site in the pro-domain (not shown). *SpzX-F, Spz1C* and *DmSpz1* do not have furin recognition sites, which are the hallmark of neurotrophins, such as *Spz2* and Spz*5* [27].

Structurally, SpzX-F/C/H have lost the conserved Cys residue involved in the formation of the intermolecular disulphide bonding (**Fig. S12d**). Instead, they have a cysteine residue in an alternative location that is predicted to form an inter-molecular disulphide bond located at the ligand-receptor interface. This residue is shared by Spz3/4/5 and Spz1C. The structure of Spz1C confirms the presence of two intermolecular disulphide bridges that constrain the covalent cystine-knot dimer doubly.

Interestingly, OrthoMCL-DB congregated *Aedes SpzX* and *Drosophila Spz1* genes into an ortholog group based on their sequence similarity (OG6_117926). *Spz1C* however is excluded from this group and forms a group on its own (OG6_426965). Next, we checked if *Spz1C* was restricted to the *A. aegypti* lineage. When this study was initiated, it seemed to have no counterpart in any of the sequenced organisms. It therefore appeared to be a lineage-restricted gene. *A. albopictus,* which diverged only ~71 Mya from *A. aegypti,* has recently been re-sequenced[28].The updated genome of *A. albopictus* contains 3 copies of *Spz1C*-like genes at loci LOC109412940, LOC109424625 and LOC115260685. Hence, *Spz1C* has undergone *Aedes*-specific duplication and a rapid expansion in *A. albopictus*, whereas *Toll1/5* sequences all appear to cluster in the same ortholog group (OG6_106857).

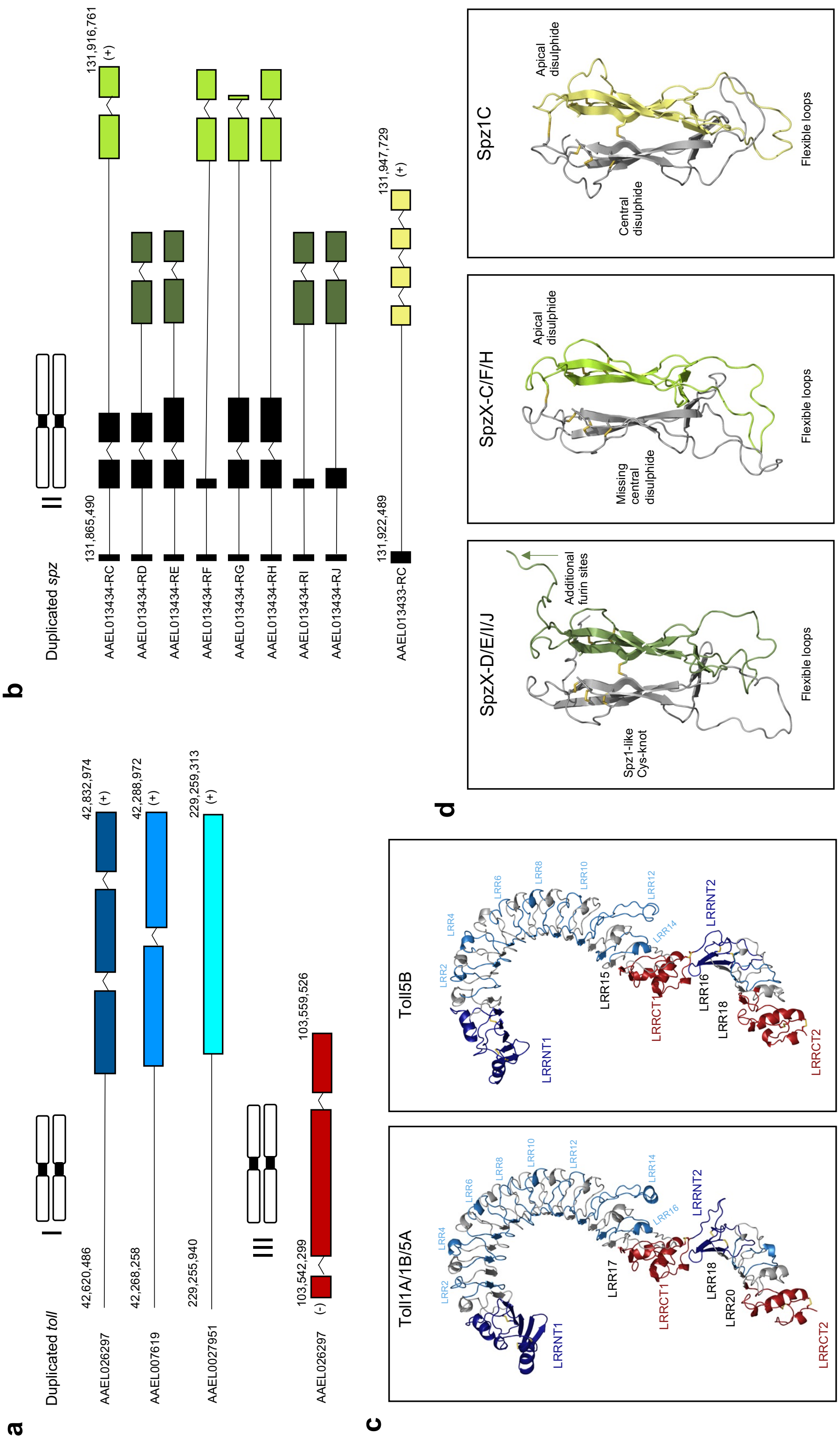

**<u>Supplementary Figure 12</u>: Similarities and differences in duplicated Toll and Spz in *Aedes aegypti*.** Gene localisation and exon/intron organisation in the Toll1/5 family **(a)** with Toll1A (AAEL026297), Toll1B (AAEL027951), and Toll5A (AAEL007619) on chromosome I, Toll5B (AAEL000057) on chromosome III. Organisation of duplicated Spz1 genes **(b)** with all family members located on chromosome II. Spz1A and Spz1B have been reannotated as mutually exclusive splice forms of a single gene labelled SpzX (AAEL003434), while Spz1C sequence and location (AAEL013433) was confirmed by high throughput sequencing techniques. **(c)** Toll ligand-binding ectodomain, and **(d)** Spz C-terminal active domain homology models generated by Modeller are visualised using PyMol.

**Supplementary Table 1: Cryo-EM Data Collection, refinement, and validation Statistics.**

| | #1 homodimer (EMDB-11984) (PDB 7B1D) | #2 heterodimer (EMDB-11982) (PDB 7B1B) | #3 heterotrimer (EMDB-11983) (PDB 7B1C) |
|---|---|---|---|
| **Data collection and processing** | | | |
| Magnification | 105,000 | | |
| Voltage (kV) | 300 | | |
| Electron exposure (e–/Å$^2$) | 51.10 | | |
| Defocus range (μm) | -2.7 to -1.0 | | |
| Pixel size (Å) | 0.83 | | |
| Symmetry imposed | - | | |
| Initial particle images (no.) | 507,268 | | |
| Final particle images (no.) | 85,810 | 40,153 | 42,866 |
| Map resolution (Å) | 3.41 4.23 | 4.23 | 3.74 |
| FSC threshold | | | |
| Map resolution range (Å) | 3.4 – 4.4 | 4.2 – 8.4 | 3.7 – 4.7 |
| **Refinement** | | | |
| Initial model used (PDB code) | 4LXR-based model | | |
| Model resolution (Å) | 2.2 | | |
| FSC threshold | | | |
| Model resolution range (Å) | 3.4 – 4.4 | 4.2 – 8.4 | 3.7 – 4.7 |
| Map sharpening *B* factor (Å$^2$) | -82.8 | -93.5 | -54.0 |
| Model composition | | | |
| Non-hydrogen atoms | 10,967 | 12,870 | 17,197 |
| Protein residues | 1,505 | 1,697 | 2,440 |
| Ligands (Glycans) | 12 | 22 | 35 |
| *B* factors (Å$^2$) | | | |
| Protein | 82.33 | 62.77 | 88.48 |
| Ligand (Glycans) | 123.03 | 165.89 | 122.04 |
| R.m.s. deviations | | | |
| Bond lengths (Å) | 0.004 | 0.005 | 0.009 |
| Bond angles (°) | 0.693 | 0.708 | 1.201 |
| Validation | | | |
| MolProbity score | 1.79 | 2.65 | 2.47 |
| Clashscore | 6.82 | 33.26 | 14.83 |
| Poor rotamers (%) | 0.29 | 0.61 | 2.17 |
| Ramachandran plot | | | |
| Favored (%) | 93.80 | 86.38 | 90.49 |
| Allowed (%) | 6.20 | 13.62 | 9.51 |
| Disallowed (%) | 0.00 | 0.00 | 0.00 |

**Supplementary Table 2: Solution scattering of Toll5A alone and in complex with Spz1C.** Synchrotron SAXS data from solutions of Toll5A ectodomain alone and in complex with Spz1C Cys-knot domain in 50 mM NaCl, 50 mM Tris-HCl, pH 7.5 were collected on the B21 beamline at the Diamond Light Source (Didcot, UK). In-line size-exclusion chromatography (SEC) SAS was employed. The SEC parameters were as follows: the samples were injected at a 0.075 ml/min flow rate onto a Superose 6 (3.2/300) column at 20°C. 31 successive 3 second frames were collected through the SEC elution peak of the sample. The data were normalized to the intensity of the transmitted beam and radially averaged; the scattering of the solvent-blank was subtracted.

| **Instrument** | **Diamond -B21** | |
|---|---|---|
| **Beam size at sample (µm)** | 1102 x 240 | |
| **Wavelength (Å)** | 0.89 – 1.3 (equivalent to 9.5 – 14 keV) | |
| ***Q* range (Å$^{-1}$)** | 0.0026 to 0.34 | |
| **Detector** | Eiger 4M (Dectris) | |
| **Detector distance (m)** | 3.7 | |
| **Exposure (s per image)** | 3 | |
| **Column** | Superose 6 Increase 3.2/300 | |
| **Flow rate (ml/min)** | 0.075 | |
| **Sample volume (µl)** | 55 | |
| **Sample concentration (mg/ml)** | 5 | |
| **Temperature (K)** | 293 | |
| | **Structural parameters** | |
| **$R_g$ (Å) Guinier** | 53.56 ± 0.05 | 55.38 ± 0.09 |
| **$R_g$ (Å) P(r)** | 55.37 | 56.62 |
| **$D_{max}$ (Å)** | 205 | 215 |
| **Porod volume (Å$^3$)** | 370,000 | 567,000 |
| | **Molecular mass determination** | |
| **Theoretical MW (kDa)** | 104 kDa (88 kDa amino acids and 16 kDa glycans) | 130 kDa ($T5A_1/S1C_1$) with S1C 26 kDa<br>234 kDa ($T5A_2/S1C_1$)<br>260 kDa ($T5A_2/S1C_2$) |
| **MALLS MW (kDa)** | 175.5 | 206.5 |
| **DATPOROD MW (kDa)** | 231 | 354 |
| **DATVC MW (kDa)** | 196 | 271 |
| **DATMOW MW (kDa)** | 202 | 267 |
| | **Data analysis software** | |
| **Data reduction** | PRIMUS & ScÅtter version 3.1r | |
| **Ab initio modelling** | DAMMIF (ATSAS version 2.8.3) | |
| **Homology modelling** | Modeller v9.24 | |
| **Computation of model fitting to data** | Pepsi-SAXS version 2.6 | |

**<u>Supplementary Table 3:</u> RT-PCR oligonucleotides used.**

| RT-PCR primers | Reference | Sequence |
|---|---|---|
| Fw Aa_Defensin_A | [74] | CTATCAGGCCGCCGTGGAG |
| Rev Aa_Defensin_A | [74] | CAATGAGCAGCACAAGCACTATC |
| Fw Aa_Dipterin_1 | [74] | GCAACATGTGGACCGATTCA |
| Rev Aa_Dipterin_1 | [74] | GTTCCTTCGTCCTGTTGATGG |
| Fw Aa_Cecropin_A | This paper | CAAAGTTATTTCTCCTGATCGCG |
| Rev Aa_Cecropin_A | This paper | CTGCACCTTCCAATTTCTTTCC |
| Fw Aa_Gambicin_1 | This paper | GTTCCTCTTGCAAGGCATATG |
| Rev_Aa_Gambicin_1 | This paper | GACAGTCACTGCAGCTTCTTATTG |
| Fw_Aa_GRRP | This paper | GCCGTTCTGGCAGTTTCCT |
| Rev Aa_GRRP | This paper | TTTTACCAACGCTACCTTGACC |
| Fw Aa_Attacin_B | This paper | TGTTCAGCGGCCAAAAGGAT |
| Rev Aa_Attacin_B | This paper | GGTTCGACGGGGTTTTGAAC |
| Fw Aa_Vago | This paper | CCTGGGAAGTGCTACGATCC |
| Rev Aa_Vago | This paper | TGGAACAGTATGCCTCGGTG |
| Fw eEF1a | [75] | AGGAATTGCGTCGTGGATAC |
| Rv eEF1a | [75] | GTTCTCTTCGGTCGACTTGC |
| Fw Aa_Toll1A | This paper | GACGTAGGTGTTCATCTTGAGG |
| Rev Aa_Toll1A | This paper | CGTACTCGCACAAGGACGA |
| Fw Aa_Toll1B | This paper | GGTCAACCTCAACGCAAACC |
| Rev Aa-Toll1B | This paper | CTTCGGGACACCTTGGTTGA |
| Fw Aa_Toll5A | [76] | TGGAAAATGATCCGATGAACTTC |
| Rev Aa_Toll5A | [76] | GTCCAGGCTGTCGATGTCTC |
| Fw Aa_Toll5B | This paper | CGACGTGTGCAGATGTAATGG |
| Rev Aa_Toll5B | This paper | TGTACCGAGCTACGCATTCC |
| Fw Aa_Toll6 | This paper | GTTGCTTTTGGTGGCGACAT |
| Rev Aa_Toll6 | This paper | ACGCATCATACAGACGGTCC |
| Fw Aa_Toll7 | This paper | GCATAGTGGAGCGCAATGTG |
| Rev Aa_Toll7 | This paper | GGCGAACGTTCCGTTTTCAA |
| Fw Aa_Toll8 | This paper | TTTCCGGACGTTGAGTAGCC |
| Rev Aa_Toll8 | This paper | CTCCAATTCCTTCAGCCCGT |
| Fw Aa_Toll9A | This paper | TATAAGGCTCCCCGTTTGCG |
| Rev Aa_Toll9A | This paper | TAGATCAGCACGGCAAGTCC |
| Fw Aa_Toll9B | This paper | GACGACTACGTCGATCACCC |
| Rev Aa_Toll9B | This paper | TCAGATTGTCCAGCAGCGTT |
| Fw Aa_Toll10 | This paper | ACATATCCCGGGACGCATTC |
| Rev Aa_Toll10 | This paper | GCGCCGGTAATGACGTAAAC |
| Fw Aa_Toll11 | This paper | GCTAACCGCATTAACGCCAG |
| Rev Aa_Toll11 | This paper | TCGTTTGACCCAAGTCGAGG |

**<u>Supplementary Source data 1</u>:** Uncropped gels relating to **Supplementary Fig. 6b** left panel: SDS-PAGE; and relating to **Supplementary Fig. 6b**, right panel: Native PAGE.

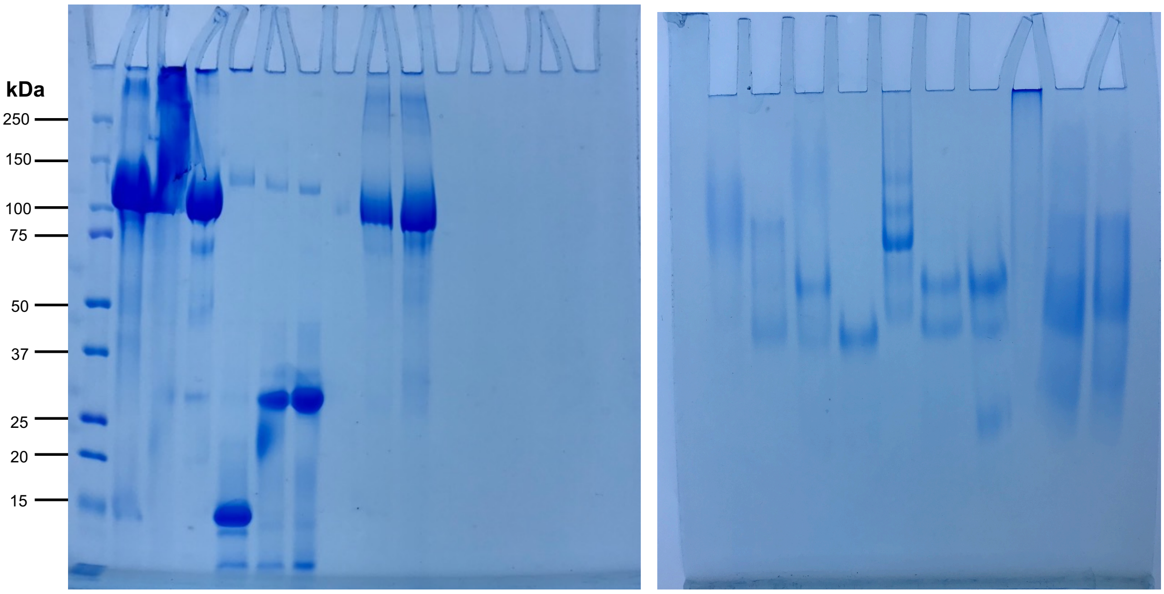

**<u>Supplementary Source data 2</u>:** Uncropped SDS-PAGE gel relating to **Supplementary Figure S7.**

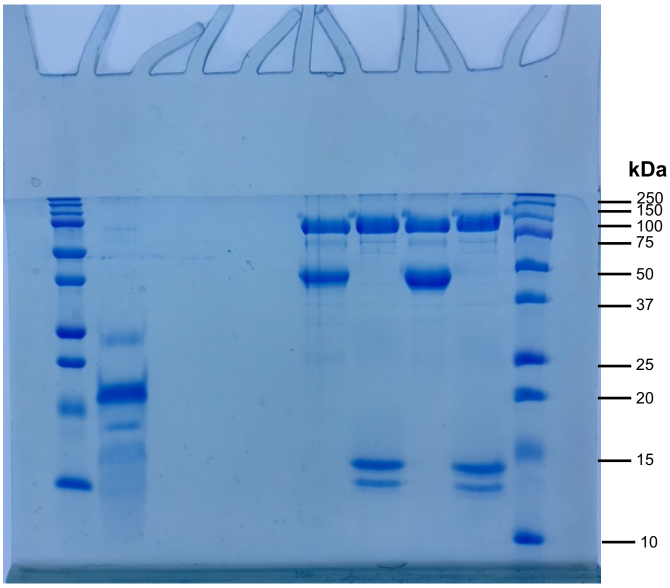

**<u>Supplementary Source data 3</u>:** Source data for agarose gels relating to **Supplementary Figure 11.**

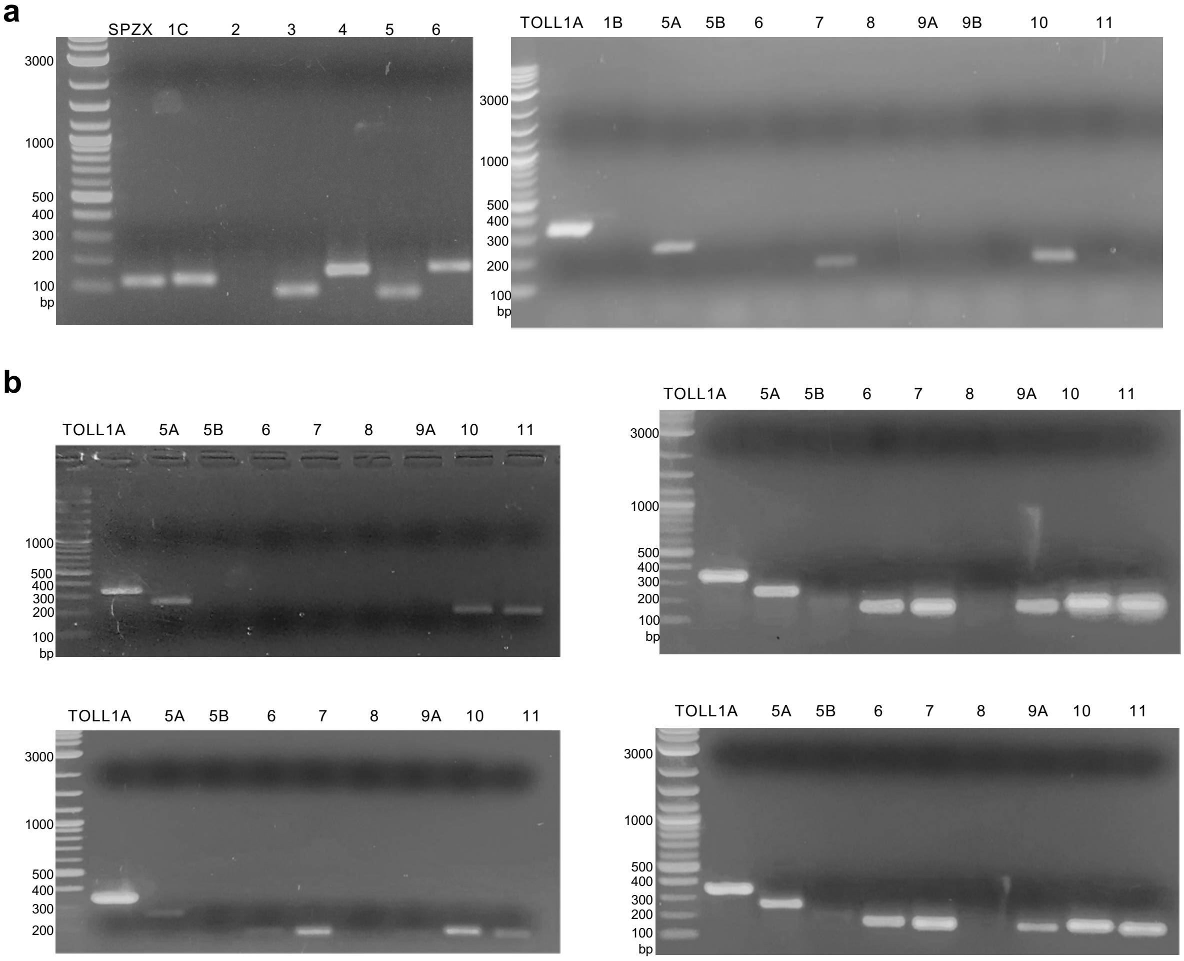
